# Enumeration, Antagonism and Enzymatic Activities of Microorganisms Isolated from Railway Station Soil

**DOI:** 10.1101/454595

**Authors:** Megha Sharma, Prashant Kaushik, Pawan Chaturvedi

## Abstract

Soil is the well-known hotspot for microbial diversity. Therefore, for our investigation, we isolated, characterized, and identified microorganisms from railway station soil. Sampling was done subsequently after every 15 days interval, and from two different soil depths i.e. 0-15 cm and below during March to May of 2013. Further, soil isolates were examined for their antagonistic activity, against four soil born plant pathogens namely, *Rhizoctonia solani, Aspergillus niger, Fusarium f sp pisi*, and *Sclerotinia sclerotiorum*. Subsequently, isolates were screened for the presence of amylases, proteases, lipases and cellulases. For each interval of soil sampling, a gradual reduction in the microbial count was noticed from month March to May. Mucor species was observed only in the rainy days. The most promising enzymes producers were *Bacillus sp., Aspergillus and Penicillium sp*. Overall, the fungal isolates were better producers of enzymes as compared to bacterial isolates.

## INTRODUCTION

Soils are the hotspots of microbial diversity and most of the soil types are rich in microbes [1]. Soil microbes interact specifically with plant root in the *Rhizosphere*, and bacterial density is generally higher in the rhizosphere [2]. Soil from places like railway station is composed of many heavy metals that is a inhibiting factor for microorganism to survive. Generally, microorganisms end up in forming separate colonies in order to produce useful enzymes like lipases, proteases, amylases and cellulases[3,4]. This enzymatic production enable them to utilize virtually every organic compound present in their niche [5].

### Material and Methods

#### Collection of sample

The study was carried from March 2013 to May 2013. Soil samples were collected regularly after every 15 days interval in clean dried ziplock bags from two different soil depths i.e. 0-15 cm and below from the Dehradun railway station, Dehradun, India.

#### Detection of colony forming unit (CFU/m^3^)

Isolation of bacteria and fungi was done by serial dilution method by using the agar plates. After incubation, the total number of colony forming unit (CFU) was enumerated in order to determine total population count for each dilution as defined elsewhere [6]. Each bacterial and fungal colony was examined carefully and purified by streak plate method on suitable agar slants. Thereafter, slants were stored at 4°C and subculturing was done after every 15 days interval.

#### Identification of bacterial and fungal cultures

The bacterial and fungal cultures were identified and differentiated based on macroscopic (shape, size, colour, margin, elevation, opacity, consistency, appearance basis of colony) and microscopic (gram staining and endospore staining) examinations [7].

#### Antagonistic test

The antagonistic activity of isolates was tested against fungal plant pathogen namely, *Rhizoctonia solani, Aspergillus niger, Fusarium f sp pisi*, and *Sclerotinia sclerotiorum*. Isolates were tested for the presence of antagonism against these plant pathogen by dual culture techniques on the potato dextrose agar plates (PDA). Further, these plates were incubated at 27±1^0^C for 5 days. There were at least three replicates of treatment and control [8]. The degree of antagonisms between each bioagent and test pathogen in dual culture was scored on scale of 1-5 as defined elsewhere [9].

### Screening of enzymatic activity

Isolates were tested by dual culture techniques and further, isolates were inoculated on different plates namely, starch agar plates (for amylase and protease), egg yolk agar plates (for lipase) and carboxymethyl cellulose agar plates (for cellulase). Further, these plates were incubated at 37°C for 24-48 hrs (for bacterial isolates) and at 27°C for 3 days (for fungal isolates) respectively. The biochemical characterization of recovered isolates were performed according to method explained elsewhere [10]. Further final proof of identity was done using the online ABIS software.

### Results

#### Total population count (CFU/m^3^) of isolates

A total of 45 bacterial and 53 fungal isolates were obtained (Table 1). The number of the bacterial colonies were decreased from March to May (Table 1). It was noticed that higher microbial count was present in upper layer of the soil i.e. 0-15 cm (Table 1).

**Table 1:** Total population count (CFU/m^3^) in soil at 2 different depth at every 15 days interval.

| Sampling dates | Bacterial Count |  | Fungal Count |  |
| --- | --- | --- | --- | --- |
|  | 0-15 cm | below 15 cm | 0-15 cm | below 15 cm |
| 04-03-2013 | 7×10<sup>9</sup> | 8×10<sup>9</sup> | 5×10<sup>9</sup> | 3×10<sup>9</sup> |
| 18-03-2013 | 7×10<sup>9</sup> | 1×10<sup>9</sup> | 3×10<sup>9</sup> | 1×10<sup>9</sup> |
| 01-04-2013 | 4×10<sup>9</sup> | 1×10<sup>9</sup> | 6×10<sup>9</sup> | 4×10<sup>9</sup> |
| 15-04-2013 | 4×10<sup>9</sup> | 1×10<sup>9</sup> | 7×10<sup>9</sup> | 5×10<sup>9</sup> |
| 29-04-2013 | 5×10<sup>9</sup> | 1×10<sup>9</sup> | 6×10<sup>9</sup> | 4×10<sup>9</sup> |
| 13-05-2013 | 3×10<sup>9</sup> | 1×10<sup>9</sup> | 4×10<sup>9</sup> | 1×10<sup>9</sup> |
| 20-05-2013 | 3×10<sup>9</sup> | 2×10<sup>9</sup> | 2×10<sup>9</sup> | 2×10<sup>9</sup> |

**Table 2:** Total occurrence of recovered isolates.

| Isolated Microorganism | Total Occurrence |
| --- | --- |
| *Bacillus subtilis* | 11 |
| *Bacillus cereus* | 7 |
| *Bacillus megaterium* | 3 |
| *Alcaligenes faecalis* | 4 |
| *Pseudomonas sp.* | 7 |
| *A. aquumarius* | 3 |
| *Staphylococcus aureus* | 6 |
| *S. equimilis* | 7 |
| *Fusarium* | 5 |
| *Aspergillus* | 8 |
| *Cladosporium* | 4 |
| *Penicillium* | 9 |
| *Epidermatophyte* | 5 |
| *Alternaria* | 7 |
| *Mucor* | 7 |
| *Rhizopus* | 7 |

#### Characterization of recovered isolates

Among the total 45 bacterial isolates identified, the frequently occurring bacteria was *Bacillus subtilis* and the least occurring were *Bacillus megaterium* and *Alcaligenes aqumarius.* However, out of 52 fungal isolates, highest count was present for the *Penicillium sp.* and the least count was determined for *Cladosporium sp.*

#### Antagonistic Test

From the isolated bacteria and fungi the isolates of *Staphylococcus aureus* was significantly positive result against all for 4 fungal plant pathogen (Table 3).

**Table 3:**
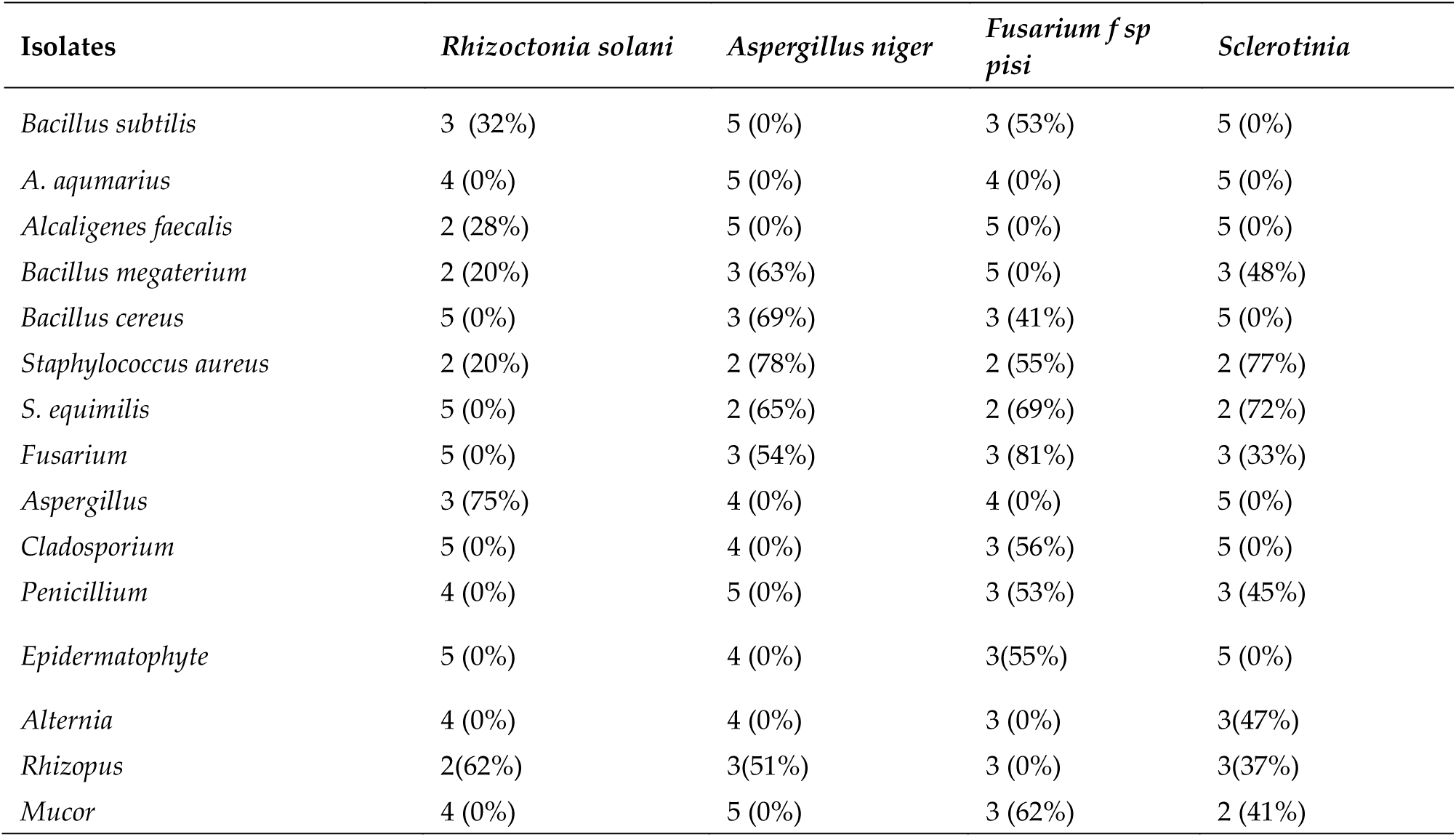
Screening of biocontrol agents against soil born fungal plant pathogens by Bell’s method and PGI %.

| Isolates | *Rhizoctonia solani* | *Aspergillus niger* | *Fusarium f sp pisi* | *Sclerotinia* |
| --- | --- | --- | --- | --- |
| *Bacillus subtilis* | 3 (32%) | 5 (0%) | 3 (53%) | 5 (0%) |
| *A. aquimarius* | 4 (0%) | 5 (0%) | 4 (0%) | 5 (0%) |
| *Alcaligenes faecalis* | 2 (28%) | 5 (0%) | 5 (0%) | 5 (0%) |
| *Bacillus megaterium* | 2 (20%) | 3 (63%) | 5 (0%) | 3 (48%) |
| *Bacillus cereus* | 5 (0%) | 3 (69%) | 3 (41%) | 5 (0%) |
| *Staphylococcus aureus* | 2 (20%) | 2 (78%) | 2 (55%) | 2 (77%) |
| *S. equimilis* | 5 (0%) | 2 (65%) | 2 (69%) | 2 (72%) |
| *Fusarium* | 5 (0%) | 3 (54%) | 3 (81%) | 3 (33%) |
| *Aspergillus* | 3 (75%) | 4 (0%) | 4 (0%) | 5 (0%) |
| *Cladosporium* | 5 (0%) | 4 (0%) | 3 (56%) | 5 (0%) |
| *Penicillium* | 4 (0%) | 5 (0%) | 3 (53%) | 3 (45%) |
| *Epidermatophyte* | 5 (0%) | 4 (0%) | 3(55%) | 5 (0%) |
| *Alternia* | 4 (0%) | 4 (0%) | 3 (0%) | 3(47%) |
| *Rhizopus* | 2(62%) | 3(51%) | 3 (0%) | 3(37%) |
| *Mucor* | 4 (0%) | 5 (0%) | 3 (62%) | 2 (41%) |

#### Enzymatic Screening of Bacterial isolates

Screening results depicted that enzyme production by different isolates varied significantly (Table 4). Generally, it was observed that none of the bacterial isolate produced cellulase. Most of the isolates produced only one kind of enzyme (Table 4). In case of fungi, *Penicillium* produced three enzymes namely, lipases, proteases and cellulases (Table 4). Generally, it was found that fungal isolates produced more enzymes as compared to bacterial isolates. Cellulases were produced by three fungal isolates namely, *Cladosporium, Penicillium, Rhizopus and Mucor*.

**Table 4:** Screening of enzymes by recovered isolates with their presence (+) and absence (-).

| Isolates | Amylases | Lipases | Proteases | Cellulases |
| --- | --- | --- | --- | --- |
| *Bacillus subtilis* | + | - | + | - |
| *Pseudomonas sp.* | + | - | - | - |
| *A. aquimarius* | + | - | - | - |
| *Alcaligenes faecalis* | + | - | - | - |
| *Bacillus megaterium* | - | - | - | - |
| *Bacillus cereus* | + | - | + | - |
| *Staphylococcus aureus* | - | + | + | - |
| *S. equimilis* | - | - | - | - |
| *Fusarium* | - | - | - | - |
| *Aspergillus* | + | + | - | - |
| *Cladosporium* | + | - | - | + |
| *Penicillium* | - | + | + | + |
| *Epidermatophyte* | - | - | - | - |
| *Alternaria* | - | - | - | - |
| *Rhizopus* | - | - | + | + |
| *Mucor* | + | - | - | + |

## Discussion

The present study gives the total bacterial count of soil samples obtained from railway station, and further indicates that the number of the bacterial colonies decreases from March to May. The presence of different microorganisms and their survival in the soil depends on the physical factors (temperature, pH, moisture content) and on the amount of organic nutrient present in the soil [11].

As noticed in our study there is increase in microbial population in 7^th^ and 9^th^ sampling due to rainy season. Recovered bacterial isolates were *Bacillus subtilis, Bacillus cereus, Bacillus megaterium, Alcaligenes faecalis, Pseudomonas sp, A. aqumarius, Staphylococcus aureus* and *S. equimilis* and fungal isolates were *Fusarium, Aspergillus, Cladosporium, Penicillium, Alternaria, Rhizopus and Mucor*. Previously, it was also shown that water is a major factor responsible for the variation of bacteria and fungi in the soil [12].

The present work on the plant pathogenic fungi showed that the many isolates can control these pathogens very efficiently. Results shows that fungal isolates like Fusarium, Rhizopus, Penicillium, Aspergillus and bacterial isolates like *Pseudomonas sp., S.aureus and S.equimilis* were highly effective against the soil-borne plant pathogens. All of these isolates showed more than 50% inhibition against all the soil-borne plant pathogens. These isolates inhibit the growth of pathogen by producing enzymes to target cell wall of fungi and mode of their action has been multifaceted including specialities like parasitism, competition, antibiosis, and/or induced resistance [13,14]. Also, plant pathogens can be controlled by various biocontrol agents, which are readily available in environment and many of these can be isolated from soil itself [15]. Control of the pathogenic fungus, therefore, would be a cheap method, and work related to multiplication and formulations of these biocontrol agents needs to be done [16].

In our study, most of the extracellular enzymes were produced by the genus *Bacillus*. Previously it was also showed that enzymes from *Bacillus* genus were significant in targeting different kinds of carbohydrates, lipids and proteins into smaller units and members of genus *Bacillus* produce several kinds of enzymes like amylases [17,18]. However, bacterial isolates were unable to produce cellulase enzymes, similar results were found elsewhere [19].

